# Fibronectin and laminin differentially affect the inflammatory environment in microphysiological systems

**DOI:** 10.64898/2026.05.13.724930

**Authors:** Margaret Radke, Christopher J. Calo, Laurel E. Hind

## Abstract

Tissue engineered constructs are increasingly used for both modeling organs and disease *in vitro* as well as for therapeutic intervention. In addition to collagen, these constructs commonly include native extracellular matrix proteins (ECM), such as fibronectin and laminin. Given the critical role of inflammatory pathways in disease and in response to implanted materials, it is important to understand the role these proteins play in regulating the inflammatory environment. Fibronectin and laminin influence neutrophil function and endothelial activation in 2D, but their regulation of the inflammatory environment in 3D engineered constructs is not clear. For this study, we used an inflammation-on-a-chip device that includes a model blood vessel surrounded by a collagen I hydrogel with fibronectin and/or laminin. We investigated the additive effects of both proteins and a range of concentrations for each protein to determine concentration dependence. Both fibronectin and laminin have concertation dependent effects on neutrophils and the endothelium. High concentrations (50 µg/mL) of fibronectin reduced neutrophil migration, while 20 µg/mL laminin reduced neutrophil extravasation and migration, potentially due to lower ICAM-1 expression by the endothelium. Interestingly, 50 µg/mL of laminin significantly disrupted endothelial vessel formation and reduced ICAM-1 and VE-cadherin expression, likely due to significant changes in the collagen architecture. The inclusion of fibronectin and laminin, even at physiological levels, results in significant effects on neutrophil behavior, endothelial vessel formation, and collagen architecture. These proteins impact the inflammatory environment and thus need to be considered when modeling diseases and designing therapeutics, especially when neutrophils or an endothelium are involved.

**Translational Impact Statement:** This work uses an inflammation-on-a-chip device to study how fibronectin and laminin impact neutrophil behavior and vascular inflammation as these proteins are commonly used in engineered constructs. We found that fibronectin impairs neutrophil migration, while laminin decreases neutrophil extravasation and migration and at higher concentrations also prevents endothelial vessel formation. Therefore, researchers should be aware that these proteins will alter the inflammatory environment when including them in engineered constructs.

## Introduction

Over the past decade, bioengineering has made significant advancements in modeling tissues and organs *in vitro* and developing tissue engineered constructs for therapeutic intervention^1,2^. As *in vitro* model complexity increases and tissue engineered materials move to the clinic, the role of immune cells in the disease pathology being modeled or causing unwanted inflammatory responses in patients is a major focus of inquiry. These systems often leverage natural hydrogels, comprised of ECM proteins, to mimic the extracellular matrix (ECM). While natural materials have many benefits—they are physiologically relevant and biocompatible—they are also complex, which can make it difficult to isolate how specific interactions with the matrix influence cell behavior. It is, therefore, critical to understand how the concentration and composition of natural ECM hydrogels influence immune responses.

Neutrophils, the most abundant innate immune cell population, are key mediators of inflammation and contribute to the pathogenesis of many diseases including cancer^3–5^, cardiovascular disease^6^, autoimmune disease^7^, and fibrosis^8–10^. During inflammatory challenges, neutrophils are recruited from blood vessels, migrate through tissue, and perform effector functions to clear pathogens and maintain tissue homeostasis^11^. Dysregulation of these processes leads to chronic inflammation and disease progression. In tissue engineered constructs, neutrophils play a major role in the foreign body response to implanted biomaterials and can drive persistent inflammation and degradation of hydrogel-based scaffolds^12,13^. Therefore, including neutrophils in models of disease and inflammation is critical to understanding the response *in vivo*. Neutrophil interactions with the vascular endothelium is pivotal in the early stages of the neutrophil response^14,15^. Therefore, understanding how neutrophils and endothelial cells alter their inflammatory behavior in response to different ECM proteins is critical in designing *in vitro* models of disease and engineering implantable materials.

Critically, both neutrophils and endothelial cells are responsive to ECM proteins. We have previously shown that collagen concentration and crosslinking affect the neutrophil response to infection in an endothelium-dependent manner^16,17^. Both properties influence neutrophil extravasation, while only collagen concentration impacts migration. While collagen is the predominant ECM protein, *in vivo*, ECM composition varies by tissue with differing concentrations of proteoglycans, glycoproteins, and fibrous proteins^18^. Adhesive proteins including fibronectin and basement membrane proteins such as laminin are often included in tissue models due to their importance in cell function and disease pathology. Prior studies have shown that fibronectin influences neutrophil motility and effector functions^19–21^ while laminin affects neutrophil effector functions and leukocyte extravasation^20,22^. However, these studies were largely conducted in 2D systems using isolated matrix proteins and lacked an endothelial layer. *In vivo*, neutrophils must first extravasate through the endothelium before migrating within a complex 3D tissue microenvironment composed of diverse ECM signals. Importantly, *in vitro*, matrix dimensionality significantly affects cell function, including migration^23^, that the presence of an endothelium significantly enhances neutrophil recruitment^24^, and that combinations of ECM proteins can alter cellular function through activation of distinct downstream signaling pathways. However, how ECM composition influences neutrophil behavior in the context of transendothelial migration and subsequent migration within the 3D tissue microenvironment remains poorly understood.

In this study, we used our inflammation-on-a-chip device to determine how the inclusion of laminin and fibronectin in a collagen hydrogel alters vascular inflammation and neutrophil responses in a physiologically relevant environment. We found the effects of fibronectin and laminin were concentration dependent with high concentrations of fibronectin leading to reduced migration in the ECM and high concentrations of laminin leading to both reduced extravasation and migration. For laminin, high concentrations also had a significant effect on the endothelial vessel leading to reduced protein expression and disorganized cell organization, potentially due to changes in the architecture of the hydrogel. Collectively, these results will inform researchers in the potential inflammatory impacts of including natural ECM proteins in their *in vitro* models and tissue engineered materials.

## Results

### The inclusion of 10 μg/mL fibronectin and/or laminin does not affect neutrophil extravasation, but does differentially impact migration

To determine the influence of fibronectin and laminin on vascular inflammation and the neutrophil response, we used an infection-on-a-chip system developed in our lab, that mimics a blood vessel surrounded by a model ECM, to study vascular responses and the human neutrophil response to infection (Figure 1A)^24^. Human umbilical vein endothelial cells (HUVECs) formed confluent lumens in 2 mg/mL collagen gels with 10 μg/mL fibronectin (FN) or laminin (LN), 5 μg/mL fibronectin and laminin (FN+LN), or no added protein (Ctrl). Neutrophil extravasation was then examined as a crucial early step in the neutrophil response to infection. To quantify extravasation, primary human neutrophils were seeded into the endothelial vessel, *Pseudomonas aeruginosa* was added to the device, and devices were imaged every hour for 8 hours (Figure S1-2). There were no significant differences in neutrophil extravasation across conditions, except between the Ctrl and FN+LN conditions at the 1-hour time point, where the former exhibited more extravasation than the latter (Figure 1B). These data indicate that the addition of fibronectin and/or laminin, at the concentrations tested, does not substantively affect neutrophil extravasation to *P. aeruginosa*.

**Figure 1:**
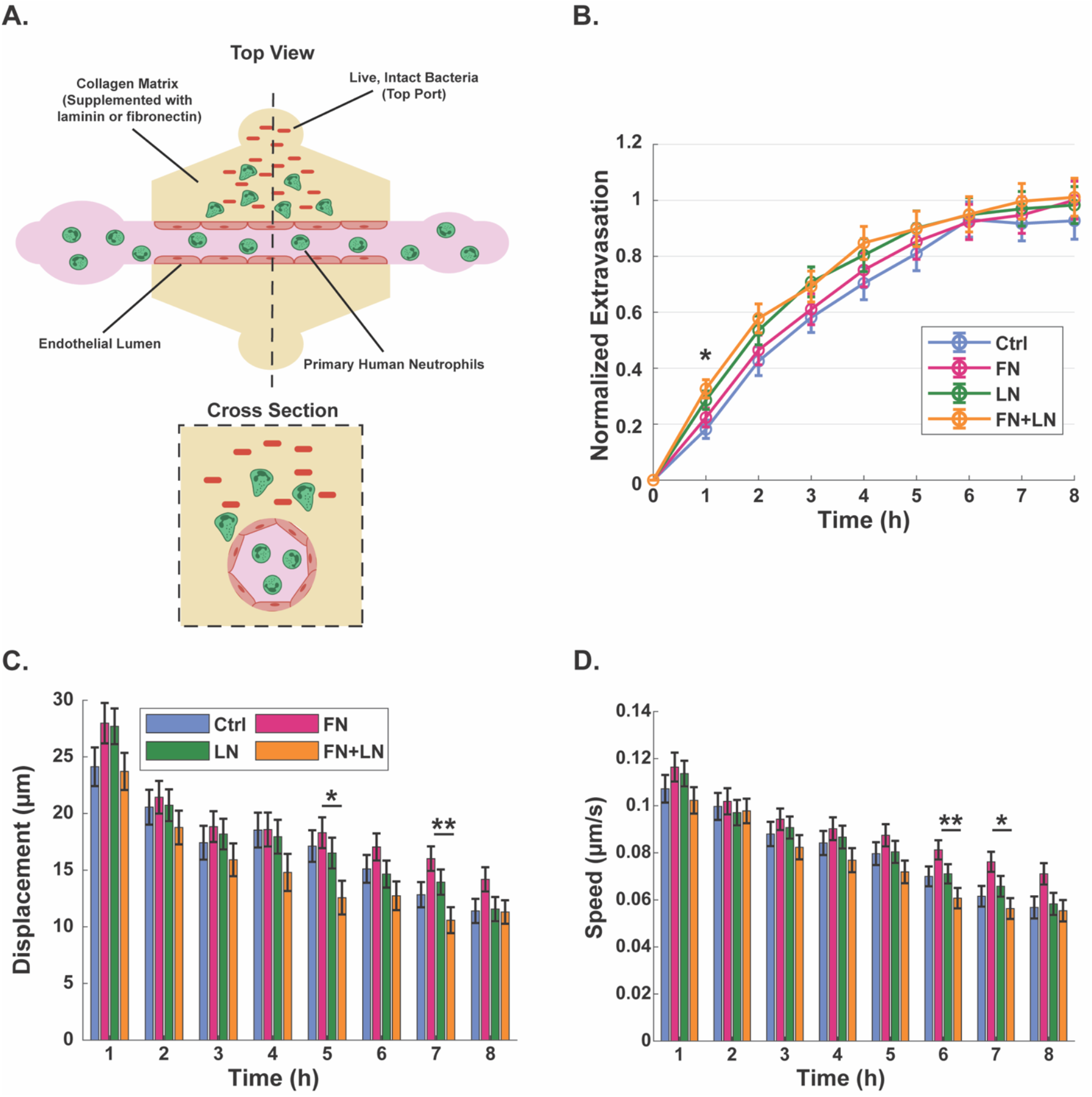
Inclusion of fibronectin and/or laminin in collagen hydrogels does not affect neutrophil extravasation, but differentially affects the migration of extravasated neutrophils. **(A)** Schematic images of the inflammation-on-a-chip device used to model a blood vessel surrounded by a model ECM comprised of 2 mg/mL collagen I hydrogels with fibronectin and/or laminin. Human umbilical vein endothelial cells (HUVECs) form an endothelial vessel, neutrophils are added inside of this vessel, and when *P. aeruginosa* is added to the top port of the device, the neutrophils extravasate and migrate towards the bacterial source. **(B)** Normalized number of extravasated neutrophils in microfluidic devices with Ctrl, fibronectin (FN), laminin (LN), and FN+LN condition matrices. Data quantified from 18 devices across 6 independent experiments and 6 neutrophil donors. Asterisks indicate significance between the Ctrl and FN+LN conditions at the same timepoint where *p<0.05. Migration properties: **(C)** displacement and **(D)** speed of neutrophils quantified over a 10-minute period every hour for 8 hours after stimulation with *P. aeruginosa*. Extravasated neutrophils were analyzed from 18 devices per condition across 6 independent experiments and 6 neutrophil donors. All conditions were compared to each other at each time point with a one-way ANOVA followed by pairwise comparisons via Tukey’s HSD test with an alpha value of 0.05. Error bars indicate the means ± SEM. Asterisks indicate significance between conditions at a given timepoint (* p<0.05 and ** p<0.01).

Once in the ECM, neutrophils must migrate through the matrix to respond to infection. To determine the effect of supplemental fibronectin and laminin in collagen on this next step of the neutrophil response, the migration properties of extravasated neutrophils in the different matrices were quantified over 10-minute intervals every hour for 8 hours post *P. aeruginosa* stimulation (Figure S3A). Neutrophils in the gels with fibronectin migrated faster and farther than neutrophils in the other conditions, with the comparison against neutrophils in the gels with fibronectin and laminin reaching statistical significance at some time points during the 5 to 7-hour timeframe (Figure 1C-D, S3B). There were no significant differences in migration straightness across conditions at any timepoint (Figure S3C). These results indicate the protein composition of the ECM differentially regulates neutrophil migration during an infectious response.

### The addition of 10 μg/mL fibronectin and/or laminin do not affect the storage modulus or pore structure of collagen gels or endothelial lumen formation

Both neutrophils and endothelial cells are sensitive to the mechanical and structural properties of the ECM^19,25–29^. Further, stiffness has been shown to impact neutrophil migration^30^. To understand if alternations in the physical properties of the ECM are leading to differences in migration, we characterized the storage moduli and pore structure of the hydrogels. To investigate this, we fabricated hydrogels with 2 mg/mL collagen I supplemented with either no additional matrix proteins (Ctrl), 10 μg/mL of fibronectin (FN), 10 μg/mL of laminin (LN), or 5 μg/mL of fibronectin and 5 μg/mL of laminin (FN+LN). Oscillatory rheology was performed on precursor gel solutions for all hydrogel conditions (Figure 2A). There were no significant differences in the storage moduli across all four conditions. After polymerization, stress relaxation tests were conducted on the gels to determine if incorporating supplemental fibronectin and laminin into collagen gels altered the viscoelastic nature of the matrices. The time it took for gels from all conditions to relax to 40% of the initial stress was not significant, suggesting no significant difference among the gels regarding stress relaxation (Figure 1B).

**Figure 2:**
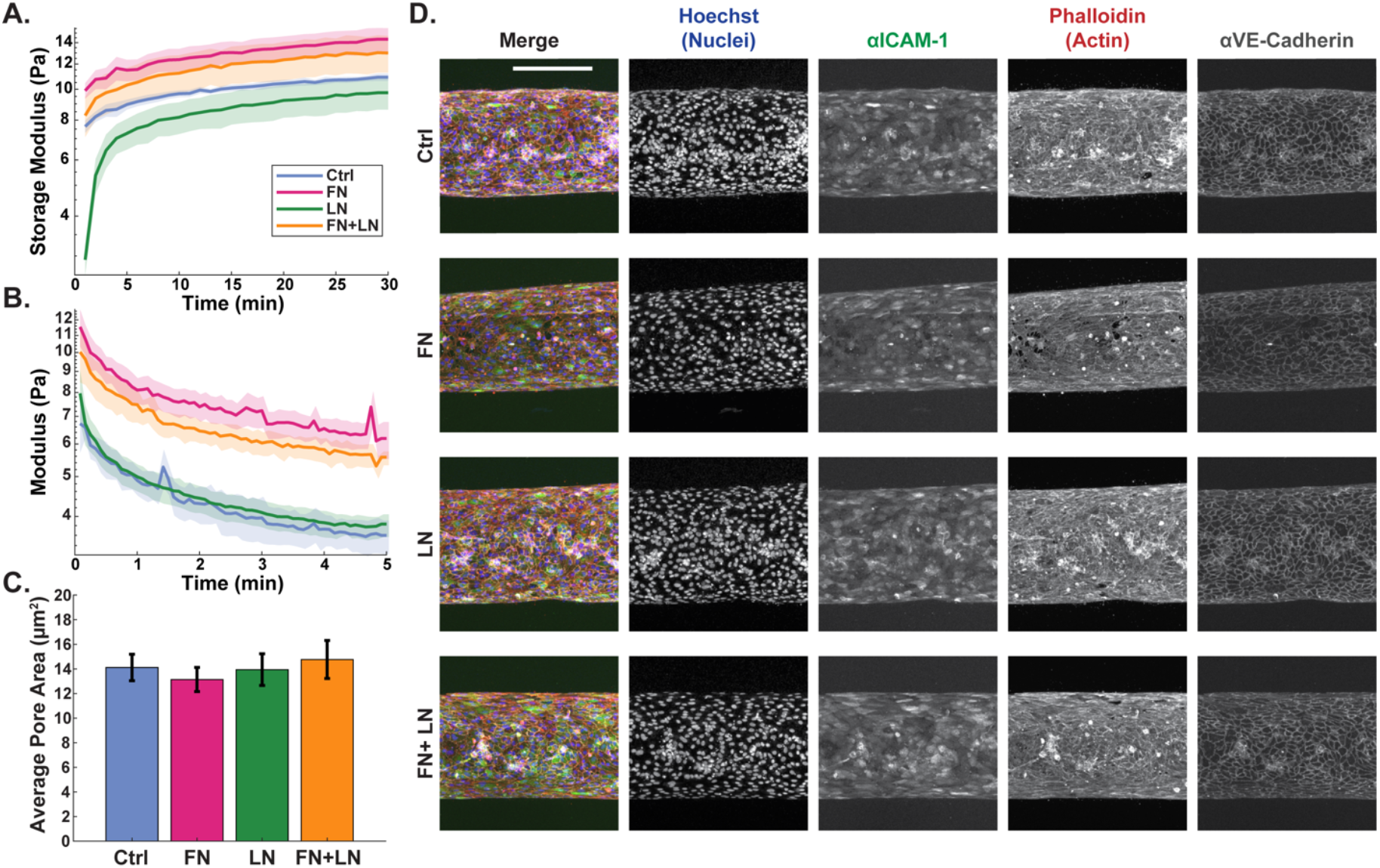
The addition of 10 μg/mL fibronectin and/or laminin do not affect the storage modulus or pore structure of collagen gels or endothelial lumen formation. Real-time rheological observations for 2 mg/mL collagen gels (Ctrl) or collagen gels supplemented with 10 μg/mL fibronectin (FN), 10 μg/mL laminin (LN), or 5 μg/mL fibronectin and 5 μg/mL laminin (FN+LN): **(A)** storage moduli (G’), 10 rad/s, 1.0% strain measured at 37 °C and **(B)** stress relaxation 15% strain at 37 °C. For A and B lines show the average modulus (n=3 gels per condition) and the shaded regions show the average modulus ± SEM. One-way ANOVA compared final modulus and 40% relaxation time across conditions. **(C)** Average area of pores for each condition. Data quantified from 4 different field of views per conditions. For pore size analysis, a Kruskal-Wallis test was performed followed by Dunn-Sidak pairwise comparisons with an alpha value of 0.05. Error bars indicate mean ± SEM. **(D)** Representative maximum intensity projections of confocal images of the bottom half of a HUVEC vessel in microfluidic devices with 2 mg/mL collagen or collagen supplemented with fibronectin and/or laminin and stained with Hoechst (nuclei, blue), anti-ICAM-1 (adhesion molecules, green), phalloidin (actin, red), and anti-VE-cadherin (tight junctions, far red) (scale bar = 250 μm).

Neutrophils largely migrate via amoeboid migration in the ECM, in which the pore size of the matrix is a major factor dictating movement^31,32^. Therefore, we quantified the pore size of the gels using confocal reflectance microscopy, where light is reflected off fibril structures to visualize collagen architecture (Figure S4A). The mean values and distributions of pore area and minor axis length (M.A.L.) were determined using Matlab’s bwlabel and regionprops functions (Figure 1C and S4B, S5). There were no significant differences in area or M.A.L. across all conditions. Together, these data show the inclusion of fibronectin and/or laminin in collagen gels at the concentrations studied do not affect the modulus or pore structure of the ECM.

As differences in neutrophil migration were not likely a result of changing matrix mechanics or collagen architecture, we next investigated if changes in the endothelial vessel might account for the change in neutrophil migration. Two days after seeding in the inflammation-on-a-chip device, HUVECs were treated with *P. aeruginosa* and stained for nuclei (Hoechst), actin (phalloidin), neutrophil adhesion molecules (ICAM-1), and tight junctions (VE-cadherin) and imaged with confocal microscopy to visualize endothelial lumen formation and differences in expression of proteins influencing extravasation in each condition. ICAM-1 facilitates neutrophil adhesion to the endothelium^33^, while VE-cadherin decreases transendothelial migration^34^, both necessary steps for neutrophil extravasation. Qualitative analysis suggests the addition of fibronectin and/or laminin does not alter endothelial vessel formation in collagen gels (Figure 2D). When quantified, the number of HUVEC nuclei is not impacted by the addition of fibronectin and/or laminin (Figure S6).

### Fibronectin alters neutrophil migration but not extravasation in a concentration-dependent manner

Previous studies have found the migration of neutrophils, along with other cell types, is impacted by fibronectin in a concentration-dependent manner^21,35,36^. Since we saw changes in neutrophil displacement and speed between conditions with 5 and 10 μg/mL of fibronectin, we wanted to explore how varying the concentration over a range of physiologically relevant values would alter the neutrophil response. For these studies 2 mg/mL collagen was supplemented with 1, 5, 10, or 50 µg/mL fibronectin.

Varying the concentration of fibronectin had no impact on neutrophil extravasation (Figure 3A). In contrast, the highest concentration of added fibronectin (50 µg/mL) reduced the speed, displacement, and path length of the extravasated neutrophils, reaching statistical significance compared to 5 µg/mL condition, 5-to 8-hours following addition of *P. aeruginosa* (Figure 3B-C, S7). Additionally, 5 µg/mL of fibronectin was identified as the optimal concentration for neutrophil migration, as cells moving through collagen with 5 µg/mL fibronectin had the maximum speed and displacement. Together, these results indicate fibronectin regulates neutrophil migration in a concentration dependent manner, with 50 µg/mL of fibronectin reducing migration, while 5 µg/mL fibronectin leads to maximum neutrophil migration without affecting the number of responding neutrophils.

**Figure 3:**
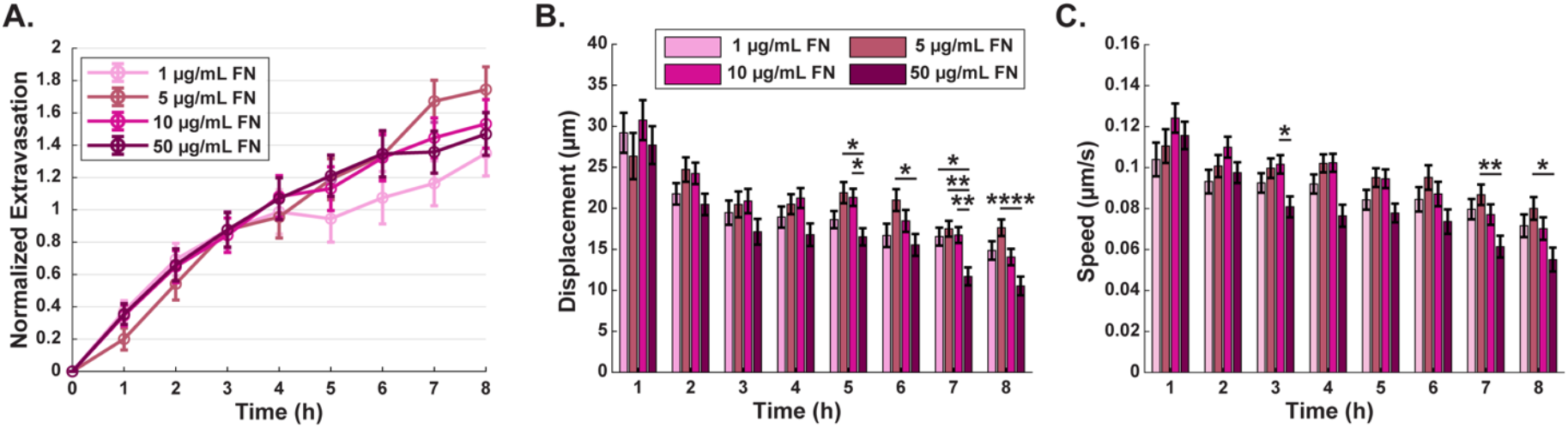
Inclusion of fibronectin alters neutrophil migration but not extravasation in a concentration-dependent manner. **(A)** Normalized number of extravasated neutrophils in microfluidic devices with 2 mg/mL collagen I hydrogels supplemented with 1, 5, 10, or 50 µg/mL fibronectin. Data quantified from 9 devices across 3 independent experiments and 3 neutrophil donors. Migration properties: **(B)** displacement and **(C)** speed of neutrophils quantified over a 10-minute period every hour for 8 hours after stimulation with *P. aeruginosa* in collagen hydrogels supplemented with 1, 5, 10, or 50 µg/mL fibronectin. Extravasated neutrophils were analyzed from 9 devices per condition across 3 independent experiments and 3 neutrophil donors. All conditions were compared to each other at each time point with a one-way ANOVA followed by pairwise comparisons via Tukey’s HSD test with an alpha value of 0.05. Error bars indicate the means ± SEM. Asterisks indicate significance between conditions at a given timepoint (* p<0.05, ** p<0.01, and **** p<0.0001).

### Inclusion of laminin reduces neutrophil extravasation and migratory properties in a concentration-dependent manner

Previous studies have found the presence of laminin 511, the isoform used in our experiments, decreases neutrophil extravasation^37^. However, we did not observe this trend in our first experiment (Figure 2B); therefore, a range of laminin concentrations (1, 5, 10, 20 µg/mL) was tested to determine if laminin’s regulation of neutrophil extravasation is concentration dependent. The highest concentration of laminin, 20 µg/mL, significantly reduced neutrophil extravasation when compared to 1 µg/mL 4 to 7-hours following the addition of *P. aeruginosa* and compared to 10 µg/mL condition at the 7-hour timepoint (Figure 4A).

**Figure 4:**
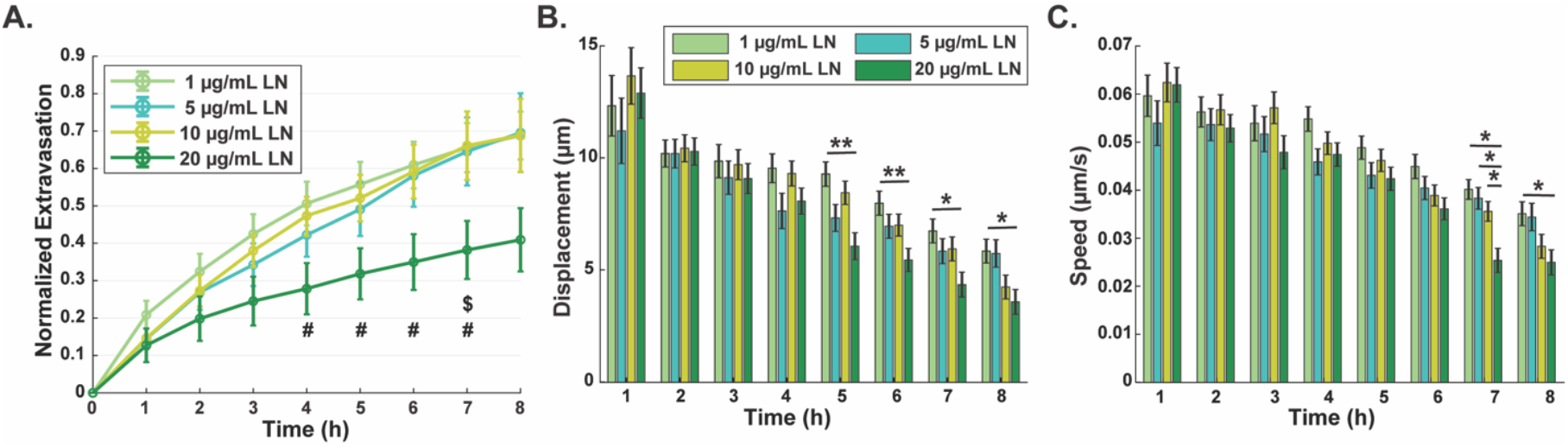
Inclusion of laminin reduces neutrophil extravasation and migratory properties in a concentration-dependent manner. **(A)** Normalized number of extravasated neutrophils in microfluidic devices with 2 mg/mL collagen I hydrogels supplemented with 1, 5, 10, or 20 µg/mL laminin. Data quantified from at least 13 devices per condition across 5 independent experiments and 5 neutrophil donors. Symbols indicate significance as # p<0.05 between the 1 and 20 µg/mL conditions, and $ p<0.05 between the 10 and 20 µg/mL conditions. Migration properties: **(B)** displacement and **(C)** speed of neutrophils quantified over a 10-minute period every hour for 8 hours after stimulation with *P. aeruginosa* in collagen hydrogels supplemented with 1, 5, 10, or 20 µg/mL laminin. Extravasated neutrophils were tracked in at least 9 devices per condition across 4 independent experiments and 4 neutrophil donors. Symbols indicate significance as * p<0.05 and ** p<0.01 between marked. All conditions were compared to each other at each time point with a one-way ANOVA followed by pairwise comparisons via Tukey’s HSD test with an alpha value of 0.05. Error bars indicate the means ± SEM.

Higher concentrations of laminin also significantly reduced neutrophil migratory properties, including speed and displacements, at later time points (Figure 4B-C). The path-length and straightness data follow the same trends (Figure S8). The maximum laminin concentration tested was 20 µg/mL, as compared to the upper concentration for fibronectin which was 50 ug/mL, because 50 µg/mL laminin results in disordered lumens while 20 µg/mL of laminin led to confluent, organized lumens (Figure 5A). The extravasation and migration data with 50 µg/mL of laminin followed the same trend as 20 µg/mL (Figure S9). These findings indicate that laminin alters neutrophil extravasation and migration in a concentration-dependent manner, with higher concentrations of laminin reducing both extravasation and migratory properties, and points to alternation in the endothelial response as one potential mechanism.

**Figure 5:**
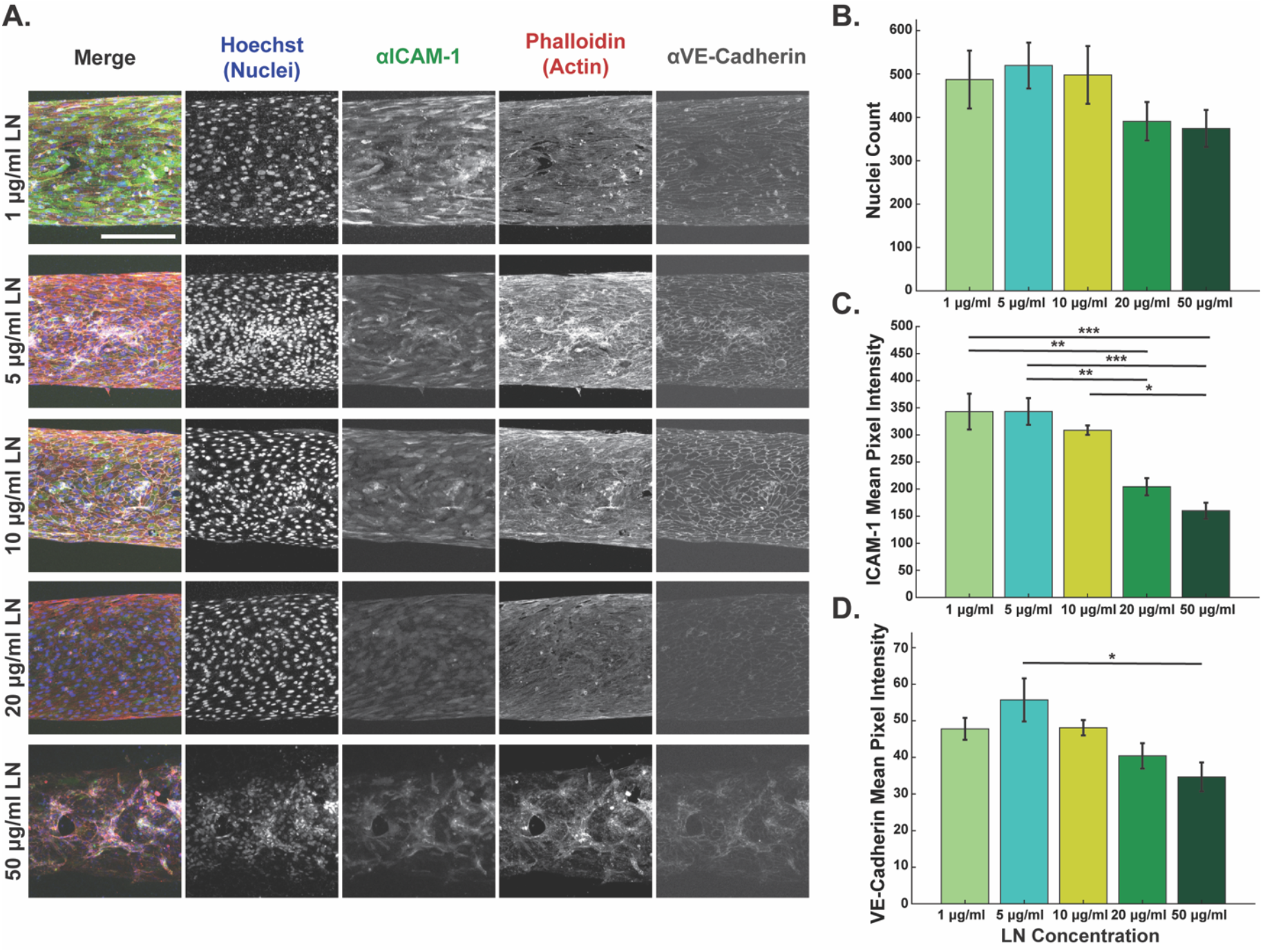
Increasing laminin concentration leads to disorganized lumens and reduced neutrophil ICAM-1 and VE-Cadherin expression. **(A)** Representative maximum intensity projections of confocal images of the bottom half of a HUVEC vessel in microfluidic devices with collagen I hydrogels supplemented with 1, 5, 10, 20, or 50 µg/mL laminin and stained with Hoechst (nuclei, blue), anti-ICAM-1 (adhesion molecules, green), phalloidin (actin, red), and anti-VE-cadherin (tight junctions, far red) (scale bar = 250 μm). **(B)** Average number of HUVEC nuclei within a set area of the lumen. Mean intensity of the stain for ICAM-1 **(C)** and VE-cadherin **(D)** within a set area. Data quantified from 47 devices across 4 independent experiments (1 µg/mL n=12, 5 µg/mL n=14, 10 µg/mL n=8, 20 µg/mL n=7, 50 µg/mL n=6). All conditions were compared to each other with a one-way ANOVA followed by pairwise comparisons via Tukey’s HSD test with an alpha value of 0.05. Error bars indicate the means ± SEM. (* p<0.05, ** p<0.01,*** p<0.001, and **** p<0.0001).

### Laminin in collagen gels leads to disorganized lumens and reduced neutrophil ICAM-1 and VE-Cadherin expression

We observed that high concentrations of laminin decreased extravasation and at 50 µg/mL laminin, endothelial organization was significantly affected; therefore, we wanted to determine the concentration-dependent effects of laminin on the endothelial lumen. Collagen was supplemented with laminin concentrations ranging from 1 to 50 µg/mL, and the endothelial lumens formed within these matrices were fixed and stained. In collagen gels with 1-20 µg/mL laminin, endothelial cells formed organized and confluent lumens, but at higher concentrations the lumens were disorganized (Figure 5A). To determine if there were differences in endothelial density or protein expression, the number of nuclei along with the intensity of ICAM-1 and VE-Cadherin were quantified from the stained images. We determined that varying the concentration of laminin does not impact the cell density of the lumens as there were no significant differences in the number of nuclei for any condition (Figure 5B). We did, however, observe changes in the protein expression. ICAM-1 and VE-Cadherin intensities decreased with increasing laminin concentration (Figure 5C and D). The reduction in ICAM-1 reached statistical significance for the 20 and 50 µg/mL conditions compared to the 1 and 5 µg/mL conditions and the 50 µg/mL condition compared to the 10 µg/mL condition. The reduction in VE-Cadherin intensity was only statistically significant between the 5 and 50 µg/mL conditions. Collectively, these results show that higher laminin concentrations significantly alter endothelial vessel formation, corresponding with a decrease in ICAM-1 and VE-Cadherin expression, when stimulated with *P. aeruginosa*.

### Inclusion of 50 µg/mL laminin increases the minor axis length and pore area of collagen gels

To understand how the highest concentrations of laminin might be disrupting endothelial vessel formation, confocal reflectance microscopy was performed. In gels with 50 µg/mL laminin, the pores are visually larger and, when quantified, the minor axis length and pore area of the gel is significantly larger (Figure 6A-C and S10). These quantified differences, along with the qualitative observation that the collagen fibrils are wider, suggests that there is more collagen bundling in the gels with 50 µg/mL laminin. These results show that laminin has a significant impact on collagen architecture, and the change in collagen pore structure could explain the high disorganized lumens observed in the gels supplemented with 50 µg/mL laminin.

**Figure 6:**
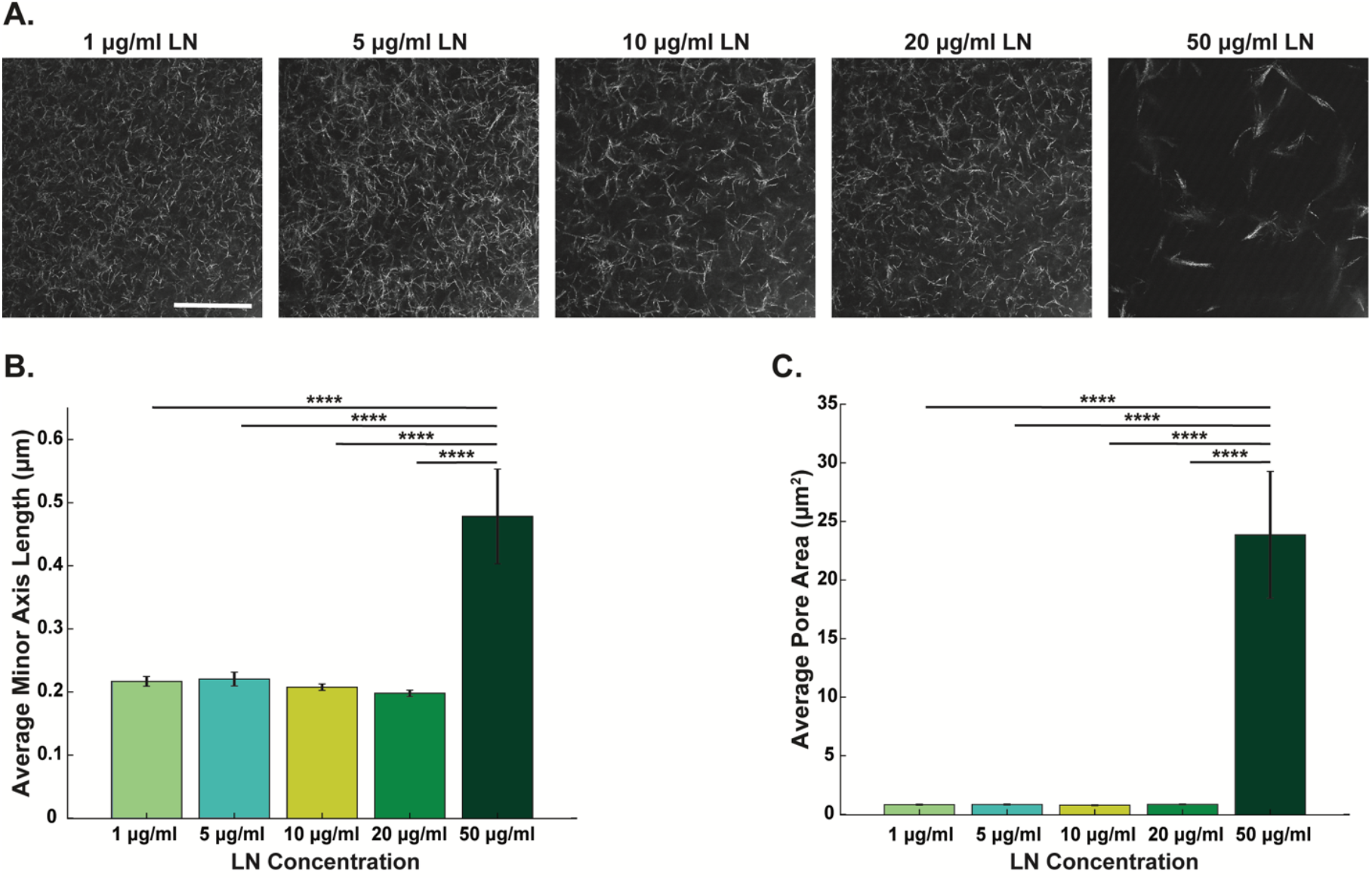
Inclusion of 50 µg/mL laminin increases the minor axis length and pore area of collagen gels. **(A)** Representative confocal reflectance images of 2 mg/mL collagen I hydrogel supplemented with 1, 5, 10, 20, or 50 µg/mL laminin (scale bar = 10 μm). Average **(B)** minor axis length and **(C)** area of pores for each condition. Data quantified from 9 gels per condition across 3 independent experiments. A Kruskal-Wallis test was performed followed by Dunn-Sidak pairwise comparisons with an alpha value of 0.05. Error bars indicate mean ± SEM (**** p<0.0001).

## Discussion

Our goal for this study was to determine the inflammatory impacts of extracellular matrix composition, specifically fibronectin and laminin, in engineered constructs. To accomplish this, we used a physiologically relevant inflammation-on-a-chip device to investigate how inclusion of these proteins in collagen hydrogels affected vascular inflammation and the neutrophil response to infection. We found fibronectin and laminin both have concentration dependent effects on the inflammatory response. High concentration (50 µg/mL) of fibronectin reduced neutrophil migration, while moderate concentration (20 µg/mL) laminin reduced both neutrophil extravasation and migration. Interestingly, increasing concentrations of laminin had significant effects on the endothelial vessels, indicating laminin may be affecting the neutrophil response indirectly through changes to the endothelium. Specifically, inclusion of 20 ug/mL laminin reduced expression of ICAM-1 compared to lower concentrations but did not affect vessel formation. In contrast, inclusion of 50 ug/mL laminin led to highly disorganized endothelial vessels and reduced expression of both ICAM-1 and VE-Cadherin, likely due to significant changes in the architecture of the collagen hydrogels. Collectively, these results highlight the potential impact of ECM protein concentration on the inflammatory microenvironment in engineered constructs.

Fibronectin is an ECM protein that is influential in cell behavior, and is important in many diseases that involve remodeling, including cancer^38,39^, fibrosis^40,41^, and biological processes such as wound healing^42,43^. Fibronectin is commonly used in both tissue modeling and implantation applications because it improves cell attachment and induces migration^35,44–46^; therefore, understanding how fibronectin impacts inflammation is important. We found fibronectin concentration had no effect on extravasation but reduced neutrophil migration at the highest concentration. Specifically, neutrophils had a biphasic response to increasing fibronectin concentration, with 5 µg/mL being an optimum concentration. This biphasic relationship in cell motility with increasing fibronectin concentration is commonly seen on 2D surfaces for a number of migrating cell types^35,47,48,36^. This was first shown in Palecek et al. using epithelial-like cells^36^. It has also been shown in chondrocytes^35^, endothelial cells^48^, and macrophages^47^ where 5 µg/mL was the optimum concentration for migration. Therefore, our finding that the relationship between motility and fibronectin concentration holds in a physiologically relevant environment is significant.

Interestingly, our finding that extravasation does not change with increasing fibronectin concentration does not fully agree with previous literature. Previous studies found fibronectin disrupts endothelial vessel formation^49^ and leads to more neutrophil transmigration^50^. Resnikoff and Schwarzbauer showed that 60 µg/mL fibronectin on a transwell insert resulted in heterogenous endothelial cell distribution and disruptions in VE-cadherin junctions^49^. Also in a transwell insert, 10 µg/mL fibronectin increased ICAM-1 expression and neutrophil transmigration compared to a coating of only collagen I^50^. Together, these studies build a narrative that fibronectin disrupts endothelial junctions and leads to more ICAM-1 expression, which both contribute to increased neutrophil transmigration, yet we did not observe this in our system. There are multiple potential explanations for why we did not see changes in neutrophil extravasation, two of which are the dimensionality of the system and the endothelial architecture. Prior studies were performed in 2D using transwells coated in fibronectin with an endothelial monolayer, whereas our system uses an endothelial vessel with fibronectin mixed throughout a collagen I hydrogel. The proteins regulating neutrophil movement are different in 2D compared to 3D^51,52^. Dimensionality is also important for the endothelium, which will change levels of protein secretion when seeded in a 2D monolayer compared to vessels^53^. Overall, the differences in dimensionality of our system compared to prior literature could explain why our results did not match literature and highlights the importance of physiologically relevant models.

Laminin is a vital protein in the basement membrane and is present throughout the ECM. Therefore, it is commonly used in tissue engineered constructs, especially in constructs used for neural tissue engineering^46,54–56^. While Matrigel is undefined, it has a high concentration of laminin, and Matrigel is used throughout biological research, making investigating how laminin impacts inflammation vital. Broadly, our neutrophil extravasation and VE-cadherin expression results matched experiments preformed *in vivo*, while the neutrophil migration^57,58^ and endothelial ICAM expression^59,60^ deviated from what would be predicted from 2D *in vitro* studies. However, the literature on how laminin impacts cells is quite inconsistent as laminin concentration, laminin isotype^22^, and cell type^61^ all contribute to laminin’s impact on cell function. Song et al., employed mouse models with various laminin isotype knockouts to investigate the effects of laminin on leukocyte extravasation. They found mice without laminin 511, the isotype used in this study, exhibited increased extravasation^22^. These results match with another study which found that *in vivo*, there is less T-cell extravasation in regions with more laminin 511^62^. These studies agree with our finding that 20 µg/mL or more of laminin reduces neutrophil extravasation. Further, Song et al. did not see changes in total VE-cadherin expression, with aligns with our findings that at laminin concentrations at or below 20 µg/mL, organized endothelial vessels form without changes in VE-cadherin expression^22^.

Conversely, our results broadly disagree with previous studies performed in 2D, and the impact of dimensionality on laminin’s effect on cells is not known. Notably, Stinson et al. found increased laminin concentration increased macrophage velocity. However, this experiment was performed in 2D, and importantly, the mechanism of migration was significantly changed by laminin concentration. At lower concentration of laminin, the cells were more spread and had a higher persistence, and with an increasing concentration of laminin, the cells were more circular. Many innate immune cells, including neutrophils and macrophages, use amoeboid migration, where cells have minimal adhesion to the ECM, and instead migrate by contracting a uropod at the trailing end of the cell^63–65^. Previous papers have shown that neutrophils migrate differently in 2D and 3D; the role of integrin regulatory proteins and receptors are different^51,52^. The changes in macrophage migration in Stinson et al. with different concentrations of laminin were attributed to changes in cell adhesion, but in 3D neutrophils migrate with minimal adhesion to the ECM, which could explain why the trends observed in this paper are not observed in our experiment. Overall, the literature on laminin’s impact on cells is convoluted, and our results generally match the *in vivo* extravasation literature, but differ from the migration literature primarily performed in 2D. This emphasizes the importance physiologically relevant geometries for studying the innate immune system, and highlights our inflammation-on-a-chip system as capable of replicating *in vivo* research.

We found high concentrations of laminin led to a disordered endothelium, possibly due to the significant change in collagen architecture we observed. These results agree with previous findings but are more severe than what has been previously seen. Grant et al. found laminin peptides were necessary for endothelial tube formation, but that high concentrations of these peptides reduces tube formation^66^. Due to the use of laminin peptides, we cannot directly compare these results to our study, but it took a much higher (400-500 µg/mL) concentration of laminin to see significant disruption in tube formation. Additionally, laminin has been shown to affect collagen architecture, with 50 µg/mL leading to a more heterogenous collagen network with punctate laminin-rich regions^67^. This study looked at architecture through scanning electron microscopy and concluded that high concentration of laminin altered the collagen network but did not see as drastic changes in the network as we observed with CRM. One reason for this could be that samples need to be dehydrated for scanning electron microscopy, which changes the structure of the collagen, while CRM does not require dehydration^68^. Overall, we found high concentration of laminin altered endothelial vessel formation and collagen architecture which aligns with what limited prior work has been performed on these topics. Despite previous research, not much is known about how these proteins affect cells in a physiologically relevant environment, as many of the previous studies were performed in 2D and if they had an endothelium, only a monolayer was used.

Broadly, the effect of including fibronectin and laminin on neutrophil and endothelial inflammation that we observed are mixed in their agreement with prior research. However, little is known about how fibronectin and laminin influence cells in a physiologically relevant environment since previous research was performed in 2D. Further, the previous studies investigating endothelial interactions used a monolayer instead of a vessel. Using an inflammation-on-a-chip system, our results closely match with *in vivo* studies, while contracting many experiments performed in 2D. This shows the importance of using engineered systems with relevant geometries and dimensionalities for investigating cell behavior.

## Materials and Methods

### Collagen preparation

High concentration type-I collagen (354249, Corning, Corning, NY) was diluted so the final hydrogel concentration was 2 mg/ml and neutralized to a pH of 7.2 using 0.5 N NaOH (02153495.5, MP Biomedicals Inc., Pittsburgh, PA), 1X phosphate buffered saline (PBS, 10010049 Gibco, Waltham, MA), and 10X PBS (BP2944100, Fisher BioReagants, Pittsburgh, PA) before adding either fibronectin (F1141-1MG, Millipore Sigma, Darmstadt, Germany), laminin (LN511, Biolamina, Sundbyberg, Sweden), or both proteins. For the fibronectin, the concentrations added were 1, 5, 10, and 50 μg/ml, while 1, 5, 10, 20, and 50 μg/mL of laminin was added. In experiments where both proteins were compared, 10 μg/mL of fibronectin or laminin, or 5 μg/mL of each protein were compared against no added protein. Solutions were kept on ice until being polymerized at 37°C.

### Rheological gel characterization

Rheology was performed on an AR 2000 ex (TA Instruments, New Castle, DE). The rheometer was fit with a Peltier Plate temperature control system and a 20 mm diameter stainless steel plate geometry. Collagen samples were prepared as previously described, and 55 ul of precursor gel solution was loaded with a 100 μm geometry gap and mineral oil around the geometry. The plate was kept at 17 °C until the start of the experiment when the temperature was raised to 37°C. Oscillatory time sweeps (10 rad/s, 1% strain) were performed for 30 minutes, then stress relaxation experiments were conducted, with 15% strain.

### Confocal reflectance microscopy

Collagen pores were visualized using confocal reflectance, as previously described^69^. Briefly, images were obtained with a confocal microscope, using a 405 nm laser and a Nikon 100x /1.45 (NA) oil immersion objective operated by Nikon Elements software.

### Pore size analysis

Using Matlab (MathWorks, 2024b), images were binarized and smoothed by non-linear filtering (mean value of neighboring pixels from a 3×3 kernel) to reduce noise. Clusters of pixels with the same values were grouped using the Matlab function bwlabel and area and minor axis length of each pore was determined using the regionprops function.

### Microfluidic device fabrication

The microfluidic devices were fabricated as previously described^16,17,70^. Additional information is included in the supplementary methods (see supplementary information).

### HUVEC culture

Pooled human umbilical vein endothelial cells (HUVEC, 50-305-964, Promocell GmbH C12203, Heidelberg, Germany) were cultured in Endothelial Growth Medium 2 (EGM-2, NC9525043, Lonza Walkersville CC3162, Basel, Switzerland). HUVECs were cultured on cell culture-treated flasks (658175, Greiner Bio-one, Monroe, NC), grown in a 37 °C incubator with 5% CO_2_ and media changes every 2 days. Cells were passaged at 80% confluency and used through passage number 6.

### *Pseudomonas aeruginosa* culture

*Pseudomonas aeruginosa* culture was performed as described previously^16,17^, and further information is included in the supplemental information.

### Device and collagen preparation

The preparation of the microfluidic device has been previously described^16,17^, and further information is included in the supplemental information.

### Stained endothelial lumens

The procedure to stain endothelial lumens has been previously described^17^ and is described in the supplemental information.

### Neutrophil isolation

All blood samples were obtained according to the institutional review board-approved protocols per the Declaration of Helsinki. Neutrophils were isolated from healthy donors’ blood, using a MACSxpress Neutrophil Isolation Kit (130-104-434, Miltenyi Biotec, Bergisch Gladbach, Germany) and a MACSxpress Erythrocyte Depletion Kit (130-098-196, Miltenyi Biotec, Bergisch Gladbach, Germany), following the manufacturer’s protocols. Informed consent was obtained from donors at the time of the blood draw according to our institutional review board.

### Neutrophil extravasation

Isolated neutrophils were resuspended in PBS with 50% of cells stained with 10 nM of calcein AM (C3100MP, Thermo Scientific, Waltham, MA), before resuspending in EGM-2 at a concentration of 7.5 million cells/mL. Approximately 5 μL of neutrophils solution were added into each lumen, followed by 3 μL of *P. aeruginosa* solution added to the top port. The devices were immediately imaged using a confocal microscope with a Nikon 10x/0.45 (NA) objective and 488 nm laser, with an environmental chamber maintaining 37 °C with 5% CO_2_ (H201-T-Unit-BL, Oko Labs, Sewickley, PA). Lumens were imaged every hour for 8 hours over a 400 μm Z-stack with 10 μm steps.

### Neutrophil migration

Neutrophils and bacteria were loaded into the device as described for extravasation experiments, but images were taken using a Nikon 20x/0.75 (NA) objective every 30 seconds for 10 minutes across a Z-stack of 80 μm with 10 μm steps every hour for 8 hours.

### Image acquisition

All imaging was performed using a Nikon A1R HD25 Laser Scanning Confocal Microscope built on the Nikon TI2-E Inverted Microscope System, a fully automated stage, and Nikon Elements acquisition software.

### Imaging processing and data analysis

Image processing has previously been described^17^, and further information is included in the supplemental information.

### Statistical analysis

For extravasation and rheology, data was pooled from three or more independent experiments, with only one value produced in a technical replicate. For migration, confocal reflectance microscopy, and endothelial staining, all the measurements within an image are averaged, and the averages of each technical replicate were pooled across the three of more independent experiments. The pore size analysis was performed with a Kruskal-Wallis test with a Dunn-Sidak correction for non-normally distributed data. All other conditions were compared to one another at each time point and analyzed with analysis of variance (ANOVA) followed by Tukey’s honestly significantly different (HSD) multiple comparison tests between conditions with an alpha value of 0.05. The mean and standard error of the mean (SEM) were calculated for each condition and P-values are labeled as *p<0.05, **p<0.01, ***p<0.001, and ****p<0.0001, unless otherwise noted.

## Conclusions

This study uses an inflammation-on-a-chip device to investigate the effects of fibronectin and laminin on inflammation. Both proteins are native to the ECM and commonly used in tissue engineered constructs, yet they both have effects on inflammatory responses. We showed concentration-dependent responses for both fibronectin and laminin on neutrophil responses and vascular inflammation. These results align with prior *in vivo* studies, but do not replicate experiments done in 2D; highlighting that system dimensionality is important considerations when designing experiments. Researchers should be cognoscente that including either of these proteins will alter the inflammatory environment, with specific changes to endothelial activation and neutrophil function but likely other immune cells as well, which can affect *in vitro* models or the immune system of the host when implanted.

## Acknowledgements

The authors would like to acknowledge and thank the Burdick Lab (namely, Morgan B. Riffe, Dr. Matthew Davidson, and Dr. Jason Burdick) at the University of Colorado – Boulder for their guidance in using their rheometer. The authors also acknowledge and thank Dr. Joseph Dragavon and the Advanced Light Microscopy Core at the University of Colorado – Boulder for his imaging expertise and maintenance of the microscopy core.

## Data Availability Statement

The raw data supporting the conclusions of this article will be made available from the corresponding author upon request.

## Ethics Statement

The studies involving humans were approved by University of Colorado Institutional Review Board, protocol number 20-0082. The studies were conducted in accordance with the local legislation and institutional requirements. The participants provided their written informed consent to participate in this study.

## Authorship

**Margaret Radke:** Formal Analysis, Investigation, Data Curation, Writing-Original Draft, Visualization, Project Administration

**Christopher J. Calo:** Conceptualization, Methodology, Software, Formal Analysis, Investigation, Data Curation, Writing-Review & Editing, Visualization, Project Administration

**Laurel Hind:** Conceptualization, Resources, Writing-Review & Editing, Supervision, Project Administration, Funding Acquisition.

