## Supplemental Data for "Fibronectin and laminin differentially affect the inflammatory environment in microphysiological systems"

**This PDF file includes:**

Figures S1 to S10

Supplementary Methods

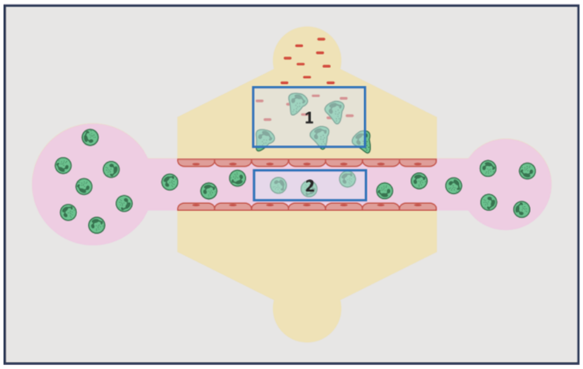


**S1: Schematic for determining normalized extravasation.** Schematic of the analysis scheme used to determine normalized extravasation. Neutrophils counted in Box 1 at all timepoints are divided by number of neutrophils counted in Box 2 at t=0 to normalize against the initial number of cells loaded into each lumen.

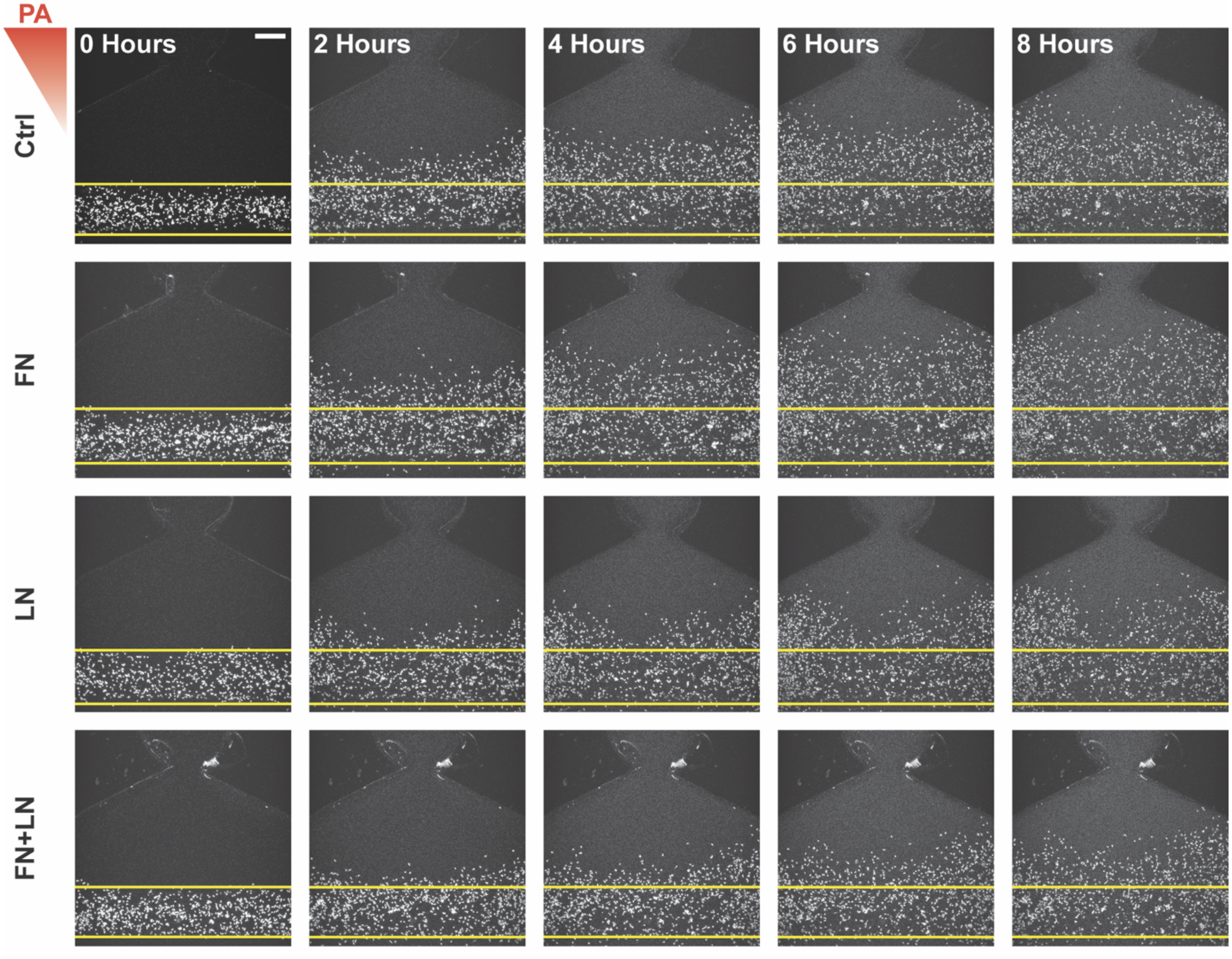


**S2: Representative images of extravasation with fibronectin and/or laminin.** Representative images of extravasated neutrophils in 2 mg/mL collagen gels (Ctrl) or collagen I gels supplemented with 10 μg/ml fibronectin (FN), 10 μg/ml laminin (LN), or 5 μg/ml fibronectin and 5 μg/ml laminin (FN+LN) at 0, 2, 4, 6, and 8 hours after stimulation with *Pseudomonas aeruginosa* (PA) (scale bar = 250 μm). Yellow lines indicate the edge of the lumen. The red triangle shows the initial bacterial gradient.

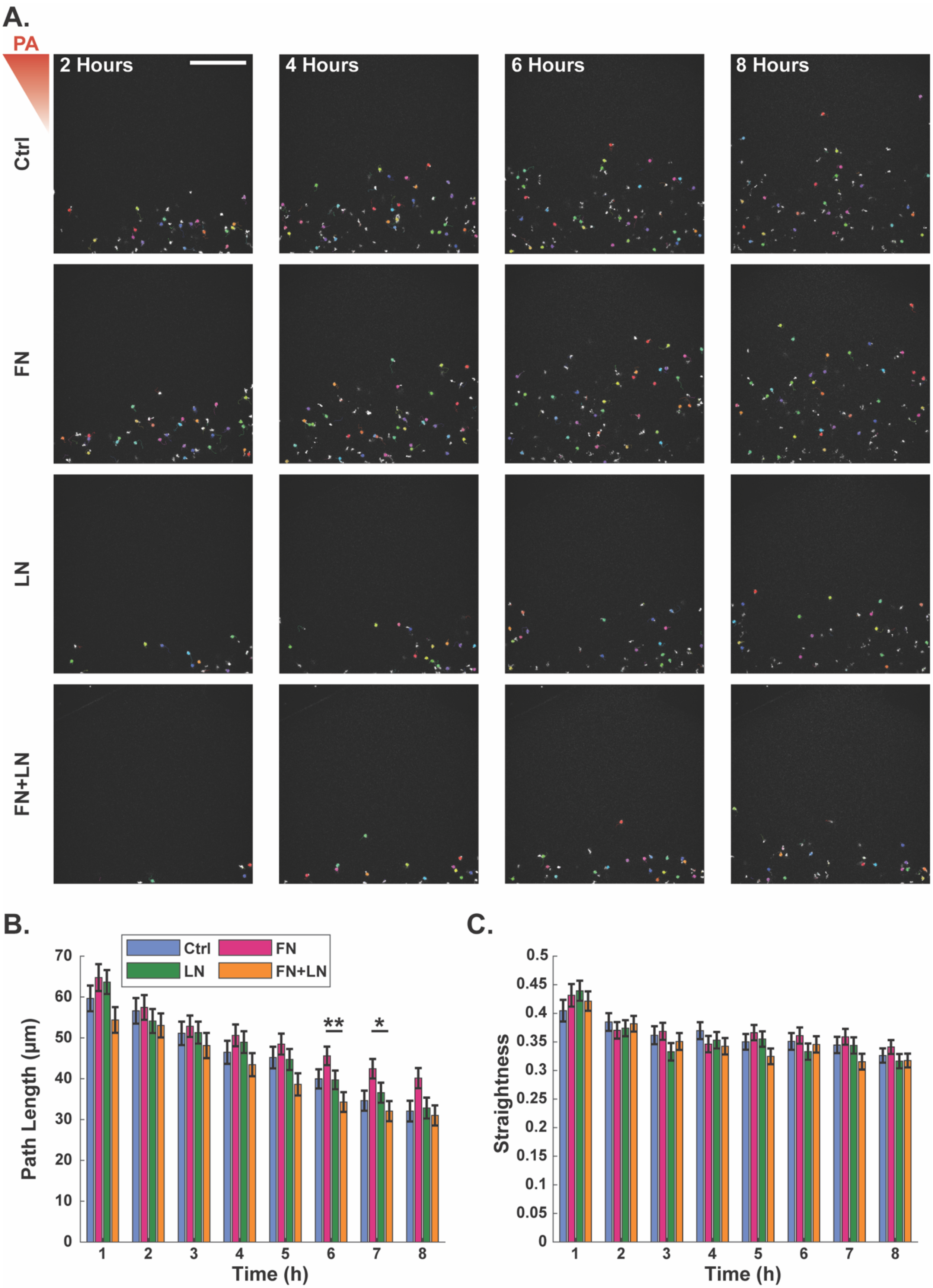
A.
PA
2 Hours
4 Hours
6 Hours
8 Hours
Ctrl
FN
LN
FN+LN
B.
Ctrl
FN
LN
FN+LN
Path Length (µm)
Time (h)
**
*
C.
Straightness
Time (h)

**S3: Supplementing collagen gels with fibronectin and/or laminin differentially affects migration of extravasated neutrophils. (A)** Representative images and tracks of migrating neutrophils in the Ctrl, FN, LN, and FN+LN condition gels at 2, 4, 6, and 8 hours after stimulation with *pseudomonas aeruginosa* (PA) (scale bar = 250 µm). Each individual track is shown in different colors. Yellow lines indicate the edge of the lumen. The red triangle shows the initial bacterial gradient. Migration properties: **(B)** pathlength and **(C)** straightness of neutrophils quantified over a 10-minute period every hour for 8 hours after stimulation with *P. aeruginosa*. Extravasated neutrophils were tracked in 18 devices per condition across 6 independent experiments and 6 neutrophil donors. All conditions were compared to each other at each time point with a one-way ANOVA followed by pairwise comparisons via Tukey's HSD test with an alpha value of 0.05. Error bars indicate the means ± SEM. Asterisks indicate significance between conditions at a given timepoint (*p<0.05 and **p<0.01).

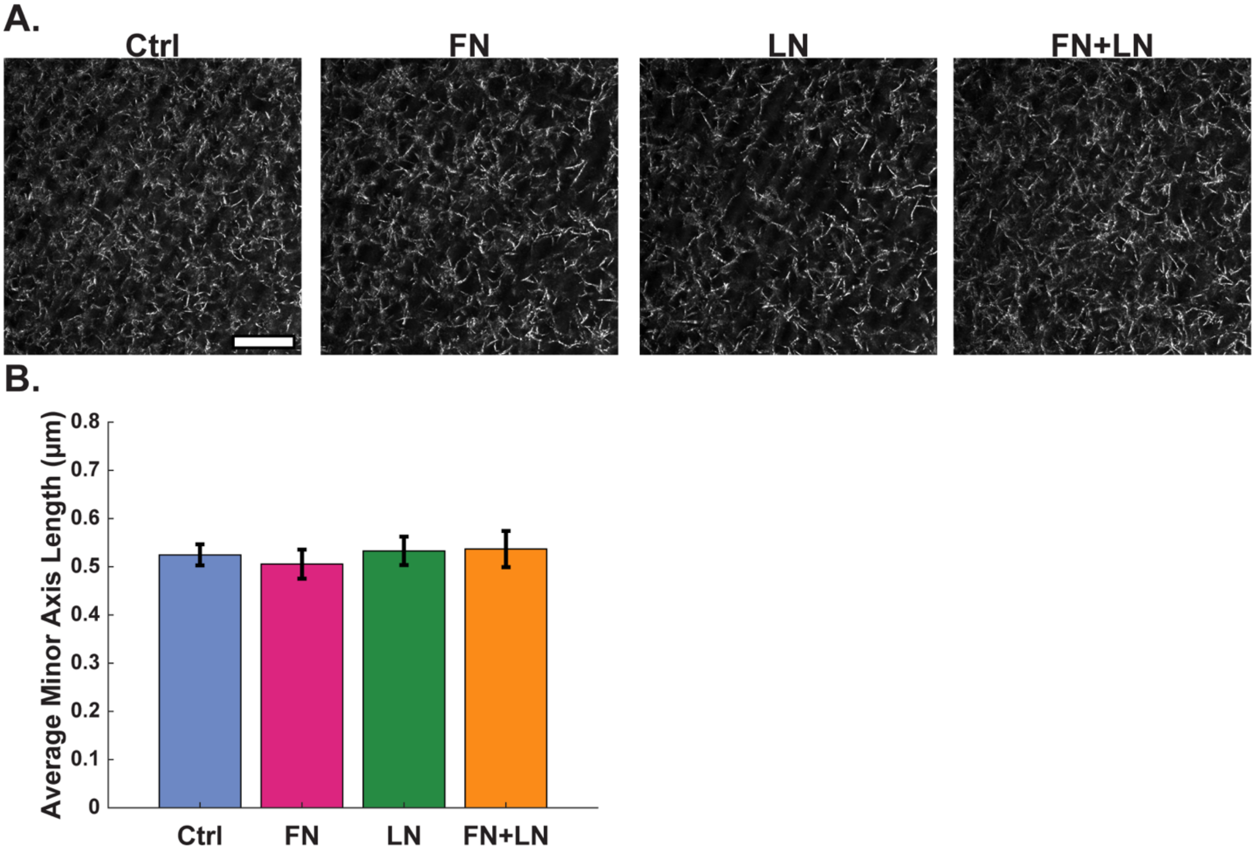


**S4: Collagen architecture is not affected by the inclusion of fibronectin and/or laminin**. (A) **(A)** Representative confocal reflectance images of Ctrl, FN, LN, and FN+LN gels (scale bar = 10 μm). **(B)** Average minor axis length of pores for each condition. Data quantified from 4 different field of views per conditions. For pore size analysis, a Kruskal-Wallis test was performed followed by Dunn-Sidak pairwise comparisons with an alpha value of 0.05. Error bars indicate mean ± SEM

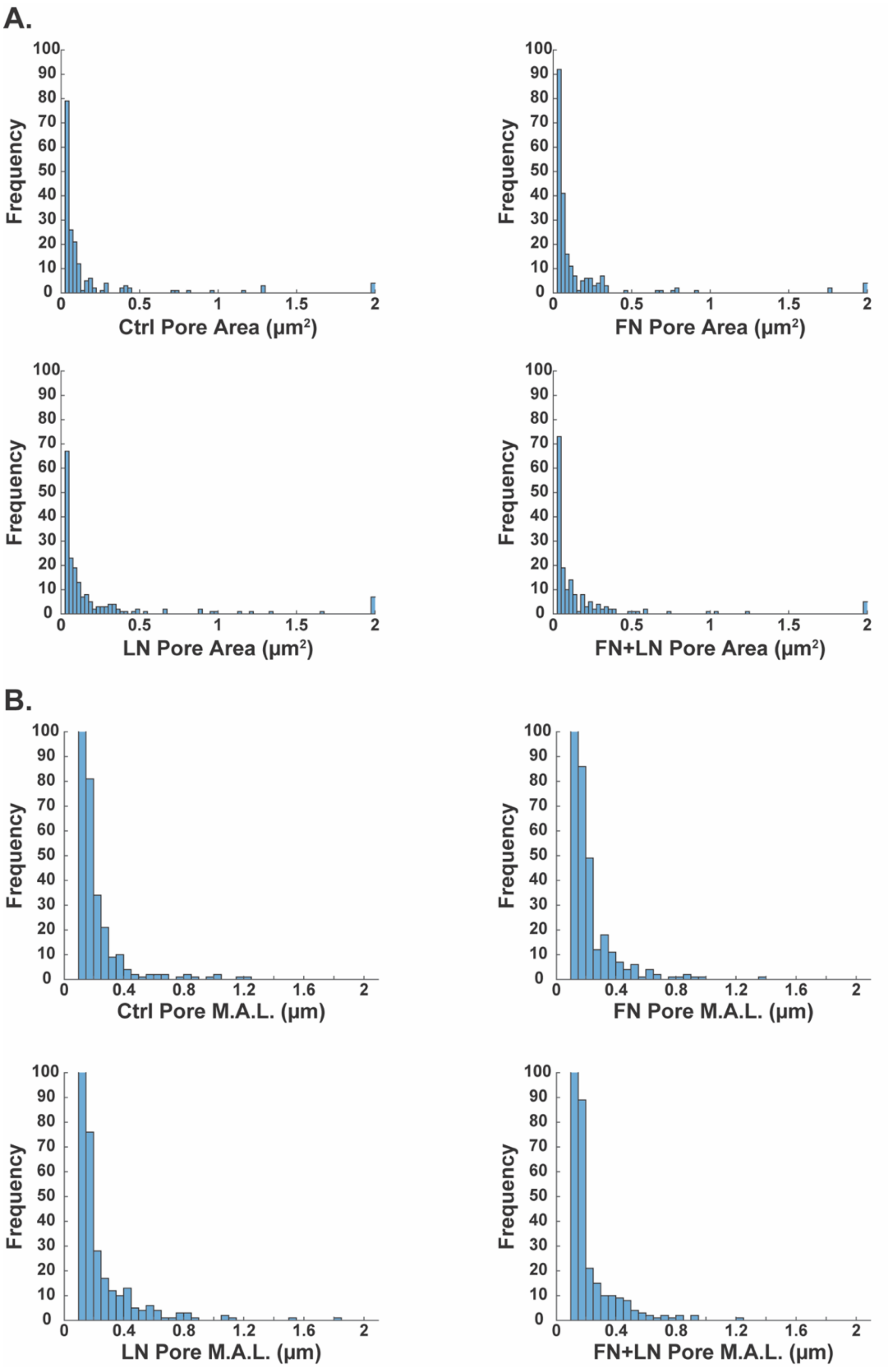


**S5: Histograms of pore area and minor axis length for collagen gels supplemented with fibronectin and laminin.** Average **(A)** area and **(B)** minor axis length (M.A.L.) for 2 mg/ml collagen gels (Ctrl) or collagen I gels supplemented with 10 µg/ml fibronectin (FN), 10 µg/ml laminin (LN), or 5 µg/ml fibronectin and 5 µg/ml laminin (FN+LN). Data quantified from 4 different field of views per conditions.

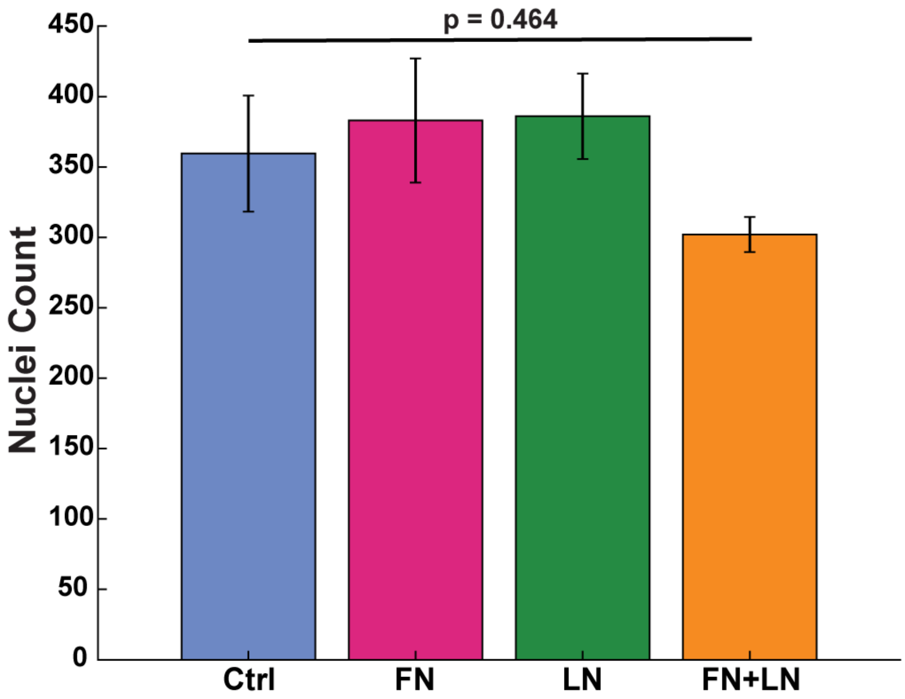


**S6: The addition of supplemental matrix proteins does not impact the number of nuclei within the endothelial vessel.** Average number of HUVEC nuclei within a set area of the lumen. Average number of HUVEC nuclei seeded in microfluidic devices with Ctrl, FN, LN, and FN+LN condition gels. Data quantified from 23 devices across 2 independent experiments (Ctrl n=6, FN n=6, LN n=7, FN+LN n=4). A one-way ANOVA test was performed.

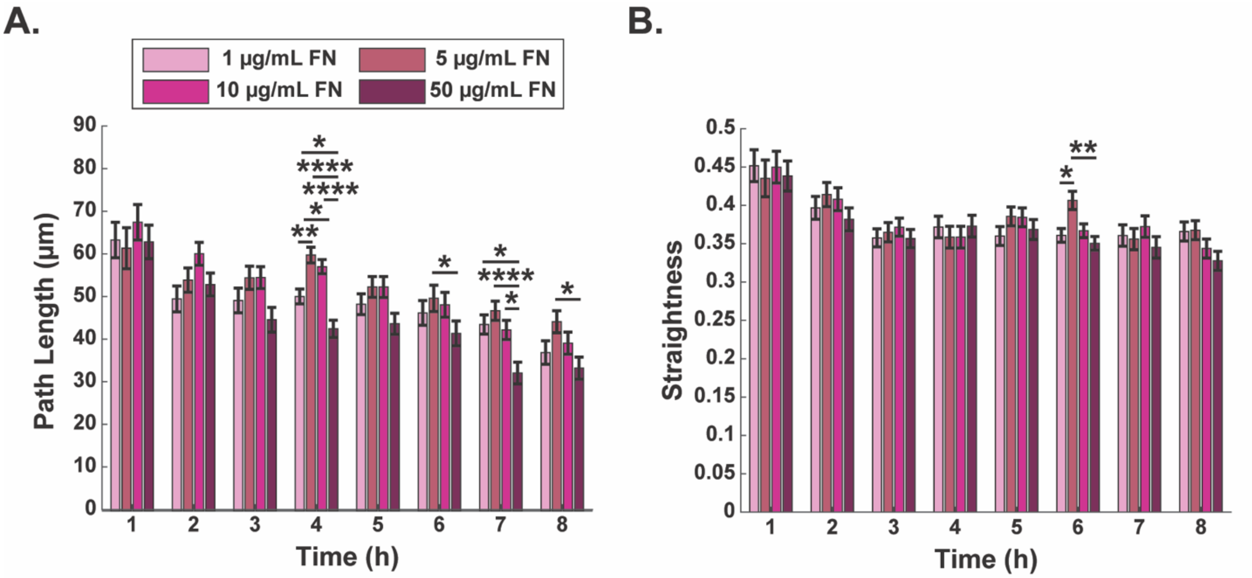


**S7: Fibronectin has a concentration dependent effect on migrating neutrophil's pathlength and a minor impact on straightness.** The **(A)** pathlength and **(B)** straightness of neutrophils quantified over a 10-minute period every hour for 8 hours after stimulation with *P. aeruginosa* in collagen I hydrogels supplemented with 1, 5, 10, or 50 µg/mL fibronectin. Extravasated neutrophils were analyzed from 9 devices per condition across 3 independent experiments and 3 neutrophil donors. All conditions were compared to each other at each time point with a one-way ANOVA followed by pairwise comparisons via Tukey's HSD test with an alpha value of 0.05. Error bars indicate the means ± SEM. Asterisks indicate significance between conditions at a given timepoint (* $p<0.05$, ** $p<0.01$, and **** $p<0.0001$).

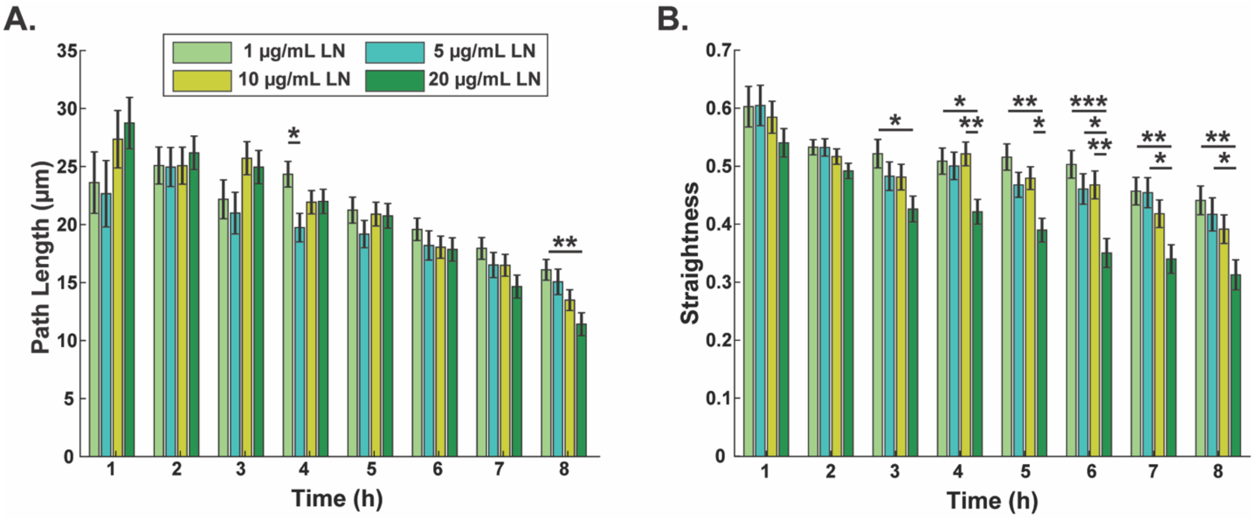


**S8: Higher concentrations of laminin reduce the migratory properties of neutrophils.** The **(A)** pathlength and **(B)** straightness of neutrophils quantified over a 10-minute period every hour for 8 hours after stimulation with *P. aeruginosa* in collagen I hydrogels supplemented with 1, 5, 10, or 20 µg/mL laminin. Data quantified from at least 9 devices per condition across 4 independent experiments and 4 neutrophil donors. All conditions were compared to each other at each time point with a one-way ANOVA followed by pairwise comparisons via Tukey's HSD test with an alpha value of 0.05. Error bars indicate the means ± SEM. Asterisks indicate significance between conditions at a given timepoint (* $p<0.05$, ** $p<0.01$, and *** $p<0.001$).

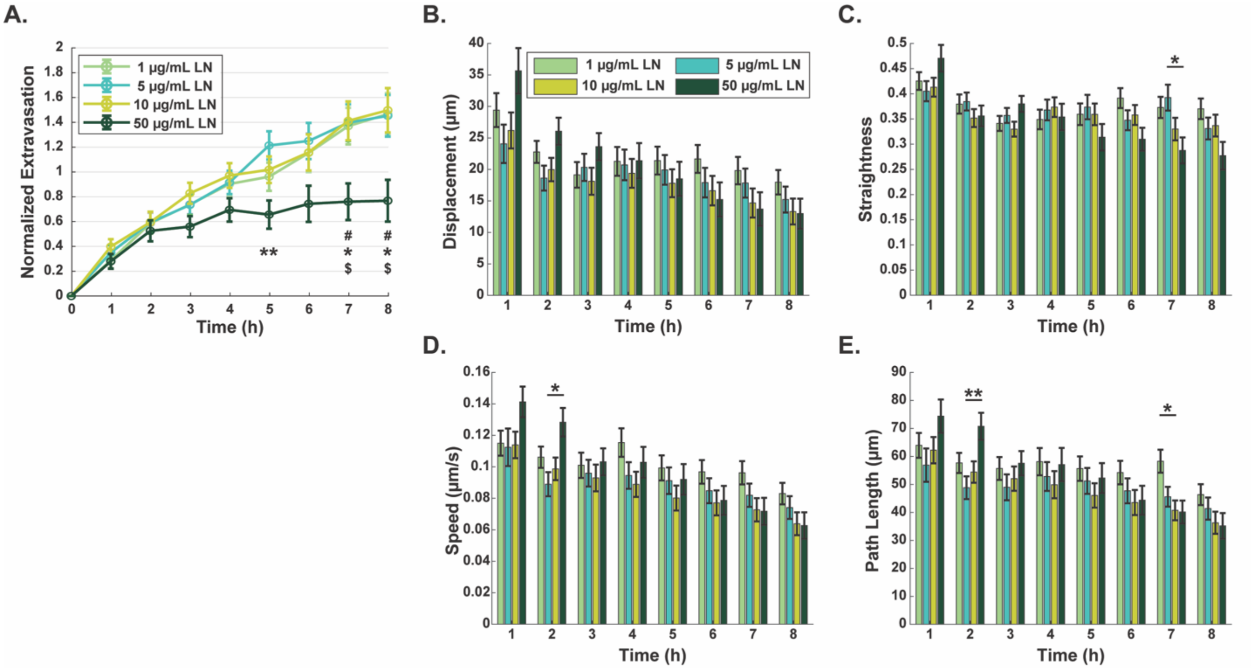


**S9: Supplementing collagen with laminin concentration up to 50 µg/mL reduces extravasation while having minimal impact on migration. (A)** Normalized number of extravasated neutrophils in microfluidic devices with 2 mg/mL collagen I hydrogels supplemented with 1, 5, 10, or 50 µg/mL laminin. Data quantified from 9 devices across 3 independent experiments and 3 neutrophil donors. Symbols indicate significance as *p<0.05 and **p<0.01 between the 5 and 50 µg/mL conditions, #p<0.05 between the 1 and 50 µg/mL conditions, and $p<0.05 between the 10 and 50 µg/mL conditions. Migratory properties: **(B)** displacement, **(C)** straightness, **(D)** speed, and **(E)** pathlength of neutrophils quantified over 10-minute increments every hour for 8 hours after stimulation with *P. aeruginosa*. Extravasated neutrophils were tracked across 9 devices per condition across 3 independent experiments and 3 neutrophil donors. All conditions were compared to each other at each time point with a one-way ANOVA followed by pairwise comparisons via Tukey's HSD test with an alpha value of 0.05. Error bars indicate the means ± SEM. Asterisks indicate significance between conditions at a given timepoint (*p<0.05 and **p<0.01).

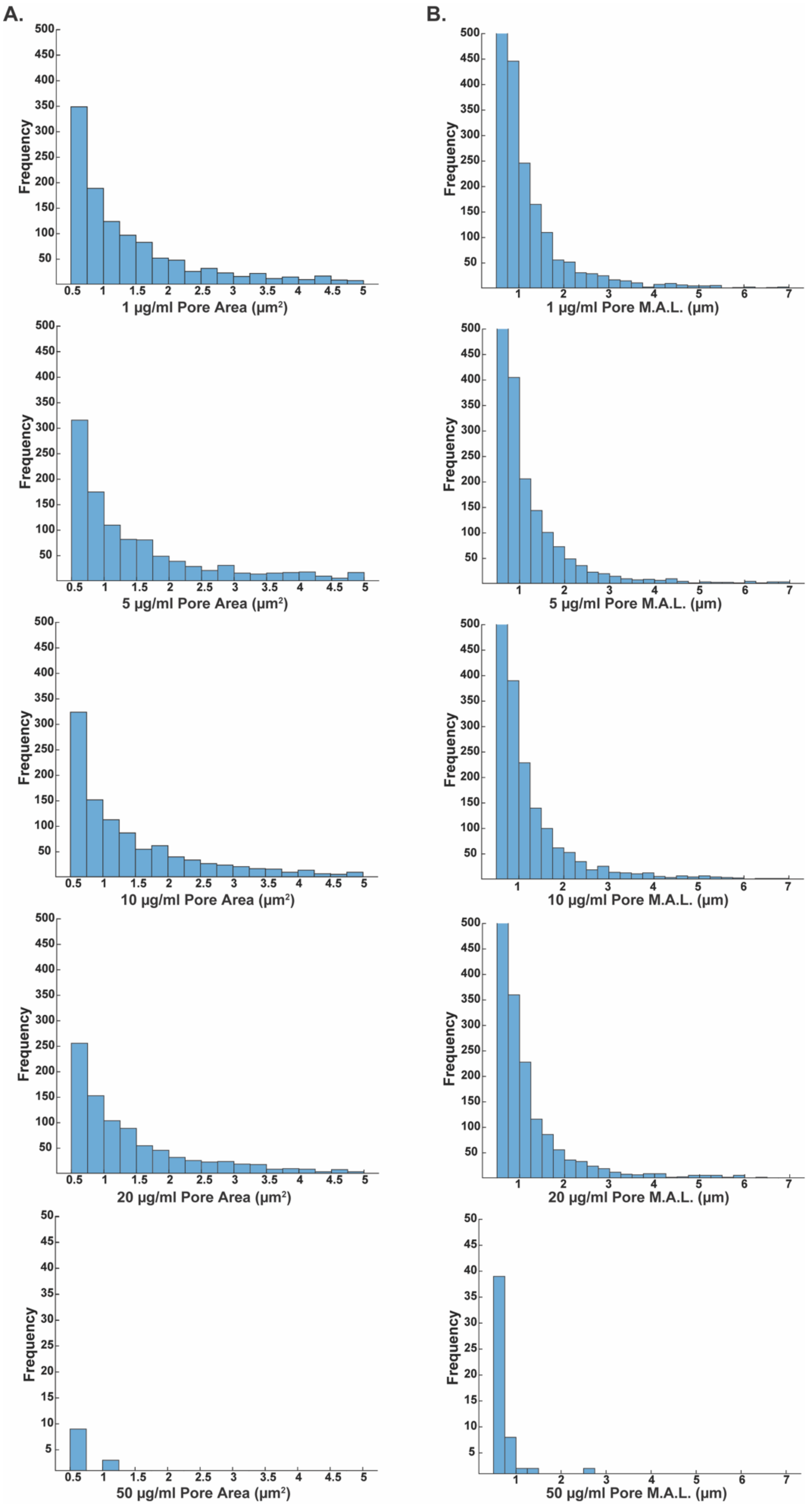

A.
B.
Frequency
1 µg/ml Pore Area (µm²)
5 µg/ml Pore Area (µm²)
10 µg/ml Pore Area (µm²)
20 µg/ml Pore Area (µm²)
50 µg/ml Pore Area (µm²)
1 µg/ml Pore M.A.L. (µm)
5 µg/ml Pore M.A.L. (µm)
10 µg/ml Pore M.A.L. (µm)
20 µg/ml Pore M.A.L. (µm)
50 µg/ml Pore M.A.L. (µm)

**S10: Histograms of pore area and minor axis length for collagen gels supplemented with varying concentrations of laminin.** Average **(A)** area and **(B)** minor axis length (M.A.L.) for 2 mg/ml collagen I gels supplemented with 1, 5, 10, 20, or 50 µg/mL laminin. Data quantified from 9 gels per condition across 3 independent experiments.

### Supplementary Methods

#### Microfluidic device fabrication

The masters for the devices were created by Protolabs Inc. using their PC-Like Advanced High Temp (Accura 5530) resin with a natural finish. Polydimethylsiloxane (PDMS, 24236-10, Electron Microscopy Sciences, Hatfield, PA) was polymerized in the molds for 4 hours at 60 ºC. The top and bottom PDMS layers were aligned, then a PDMS rod (0.337 mm inner diameter) was inserted between the layers, and the PDMS was attached to a glass-bottom dish (P50G-1.5-30-F, MatTek Corporation, Ashland, MA) with oxygen plasma from a PE25-JW Plasma Etcher (Plasma Etch, Carson City, NV).

#### *Pseudomonas aeruginosa* culture

LB plates were made according to the manufacturer's instructions (22700025, Thermo Fisher Scientific, Waltham, MA). *Pseudomonas aeruginosa* strain K (PAK) was streaked onto the plates and incubated at 37 ºC for 16 hours, then stored at 4 ºC until use. A colony was grown overnight in 5 mL of LB broth in a 37 ºC bacterial shaker. The next day, 1 mL of the cultured solution was diluted in 4 mL of fresh LB broth before being put back in the 37 ºC shaker for 1.5 hours. 1 mL of bacterial culture was pelleted using centrifugation (17,000xg for 1 minute) and resuspended in 100 µL of EGM-2. The optical density (OD) was measured at 600 nm, and the bacterial solution was diluted in EGM-2 to an OD of 5 ($1.25 \times 10^6$ CFU/mL).

#### Device and collagen preparation

The devices were UV sterilized, and the central chamber was incubated with 2% polyethylenimine solution (03880, Millipore Sigma, Darmstadt, Germany) for 10 minutes, followed by 0.4% glutaraldehyde solution (G6257, Millipore Sigma, Darmstadt, Germany) for 30 minutes. The collagen solution was prepared as described in the methods and added to the device, before allowing the collagen to polymerize overnight at 37 ºC. The PDMS rod was then removed and 5 uL of a 200,000 HUVEC/µL solution was added to this void. Devices were placed in an incubator and flipped upside down every 15 minutes for 1 hour, before changing the media. Cells were grown for 2 days with two media changes per day.

#### Stained endothelial lumens

HUVEC lumens were cultured for 2 days in the microfluidic devices, then treated with *P. aeruginosa* for 2 hours before being incubated with prewarmed 4% paraformaldehyde (PFA, AAJ19943K2, Thermo Scientific, Waltham, MA) in phosphate buffered saline (PBS, B2944-100, Thermo Scientific, Waltham, MA) for 30 minutes at 37ºC. The PFA was aspirated, followed by 3 washes with PBS, before adding PBST (PBS with 0.1% Tween 20) and incubating at room temperature for 10 minutes. Each lumen was then incubated with 33 uL of the stain solution, containing Hoescht (1:200, 23491-45-4, Sigma Aldrich, St. Louis, MO), phalloidin (1:500, ab176757, abcam, Cambridge, United Kingdom), anti-ICAM (1:200, BBA20, R&D Systems, Minneapolis, MN), and anti-VE-cadherin (1:120, 561567, BD Pharmingen, Franklin Lakes, NJ) in PBST overnight at 4 ºC. The next day, the center channel of the device was washed 3 times with PBS to remove any excess stain before being stored with PBS in the lumen at 4ºC before imaging. Imaging was preformed using a confocal microscope with a Nikon 20x/0.95 (NA) water immersion objective operated by Nikon Elements software, with 405 nm, 488 nm, 561 nm, and 640 nm laser lines. Lumens were imaged with a 150 μm Z-stacks with 2 μm steps along the Z-axis.

**Image processing and data analysis**

Extravasation, migration, and endothelial staining experiments were analyzed in the Nikon Elements software. Max intensity projections in the Z-axis were created for each file. For extravasation and migration experiments, only neutrophils from the center of the device were analyzed. For extravasation, a 900 μm x 600 μm rectangular region of interest resting above the top of the endothelial vessel centered on the device was created. The neutrophils within the region were counted at each time point using the bright spots function in the 488 nm channel within Nikon Elements Analysis Software. The number of neutrophils extravasated was normalized to the initial number of neutrophils to account for any differences in cell loading (Figure S1). The initial number of neutrophils is counted within a 900 μm x 300 μm rectangular region in the middle of the lumen from the first time point.

For migration experiments, neutrophils were tracked in the 488 nm channel using the cell motility function within Nikon Elements Analysis Software. Cells were identified through bright spot detection, and cells that remained in frame through at least half of the images were recorded. Net displacement was calculated as the distance between the starting and ending positions for each interval. Migration path length was output by the software for each neutrophil and was used with displacement to determine the straightness of migration: the path length of a neutrophil divided by its net displacement. Lastly, speed was computed as the total path length travelled by a neutrophil divided by the time elapsed during the track.

Quantification of the stained endothelium was performed within the Nikon Elements Analysis Software. The analysis is performed within a 550 μm x 350 μm region placed in the center of the endothelial lumen. Nuclei were identified by thresholding the Hoechst images and counted using the bright spot function. Within the region, the average intensity of the anti-ICAM and anti-VE-Cadherin-stained images was calculated.
